# GRIP: Graph Representation of Immune Repertoire Using Graph Neural Network and Transformer

**DOI:** 10.1101/2023.01.12.523879

**Authors:** Yongju Lee, Hyunho Lee, Kyoungseob Shin, Sunghoon Kwon

**Author notes:** These authors contributed equally (Yongju Lee, Hyunho Lee, Kyoungseob Shin).

## Abstract

The immune repertoire is a collection of immune receptors that has emerged as an important biomarker for both the diagnostic and therapeutic of cancer patients. In terms of deep learning, analyzing immune repertoire is a challenging multiple-instance learning problem in which the immune repertoire of an individual is a bag, and the immune receptor is an instance. Although several deep learning methods for immune repertoire analysis are introduced, they consider the immune repertoire as a set-like structure that doesn’t take into account the nature of the immune response. When the immune response occurs, mutations are introduced to the immune receptor sequence sequentially to optimize the immune response against the pathogens that enter our body. As a result, immune receptors for the specific pathogen have the lineage of evolution; thus, the immune repertoire is better represented as a graph-like structure. In this work, we present our novel method, graph representation of immune repertoire (GRIP), which analyzes the immune repertoire as a hierarchical graph structure and utilize the collection of graph neural network followed by graph pooling and transformer to efficiently represents the immune repertoire as an embedding vector. We show that GRIP predicts the survival probability of cancer patients better than the set-based methods, and graph-based structure is critical for performance. Also, GRIP provides interpretable results, which prove that GRIP adequately uses the prognosis-related immune receptor and gives the further possibility to use the GRIP as the novel biomarker searching tool.

## Introduction

The importance of the immune system in tumor studies is increasing because of the emerging next-generation cancer therapies that directly target immune cells or utilize them as mediators (Thorsson et al. 2018). The immune receptor is a protein that is expressed on the surface of the immune cell to recognize the disease agents or pathogens that enter our body. It consists of an amino acid sequence that is 300 long and is important for the immune response (Slamon et al. 2011). In the case of tumors, tumor cells are pathogens in our body that immune cells recognize and attempt to defeat by infiltrating the tumor mass. Owing to the importance and utility of immune receptors, several deep learning methods have been developed to predict the properties of the individual immune receptor, such as their binding ability with specific pathogens, protein structure, and the cancer-relatedness of immune receptor (Beshnova et al. 2020; Davidsen et al. 2019; Jin et al. 2021; Mason et al. 2021)

Still, a method for analyzing the immune repertoire, a collection of immune receptors, is necessary. The role of immune receptors during the immune response can be better understood by investigating the immune repertoire. The immune repertoire shows the systematic immune status of an individual through the chronological history of the immune receptor. Thus, we can understand which immune status that immune receptor is in during the immune response. However, analyzing the immune repertoire is a challenging multiple-instance learning (MIL) problem. Although the immune repertoire consists of a few thousand to millions of immune receptors, few subsets of the immune receptor are relevant to the specific disease. Thus, to diagnose a disease based on immune repertoire, the model needs to detect the few subsets of the immune receptors from the immune repertoire (Emerson et al. 2017; Mora and Walczak 2019). In addition, even if the immune repertoire datasets of certain diseases are collated, it is difficult to recognize shared immune receptors because each individual has a unique pathogen exposure history (Greiff et al. 2017). Previous deep learning methods attempted to solve the challenging problem of set-based MIL methods, which consider each immune receptor as an unordered (permutation invariant) dataset. All the extant methodologies, such as k-mer-based feature count or LSTM-like feature extraction combined with an attention-based module, consider the immune repertoire as a set (Figure 1a-b).

**Figure 1:**
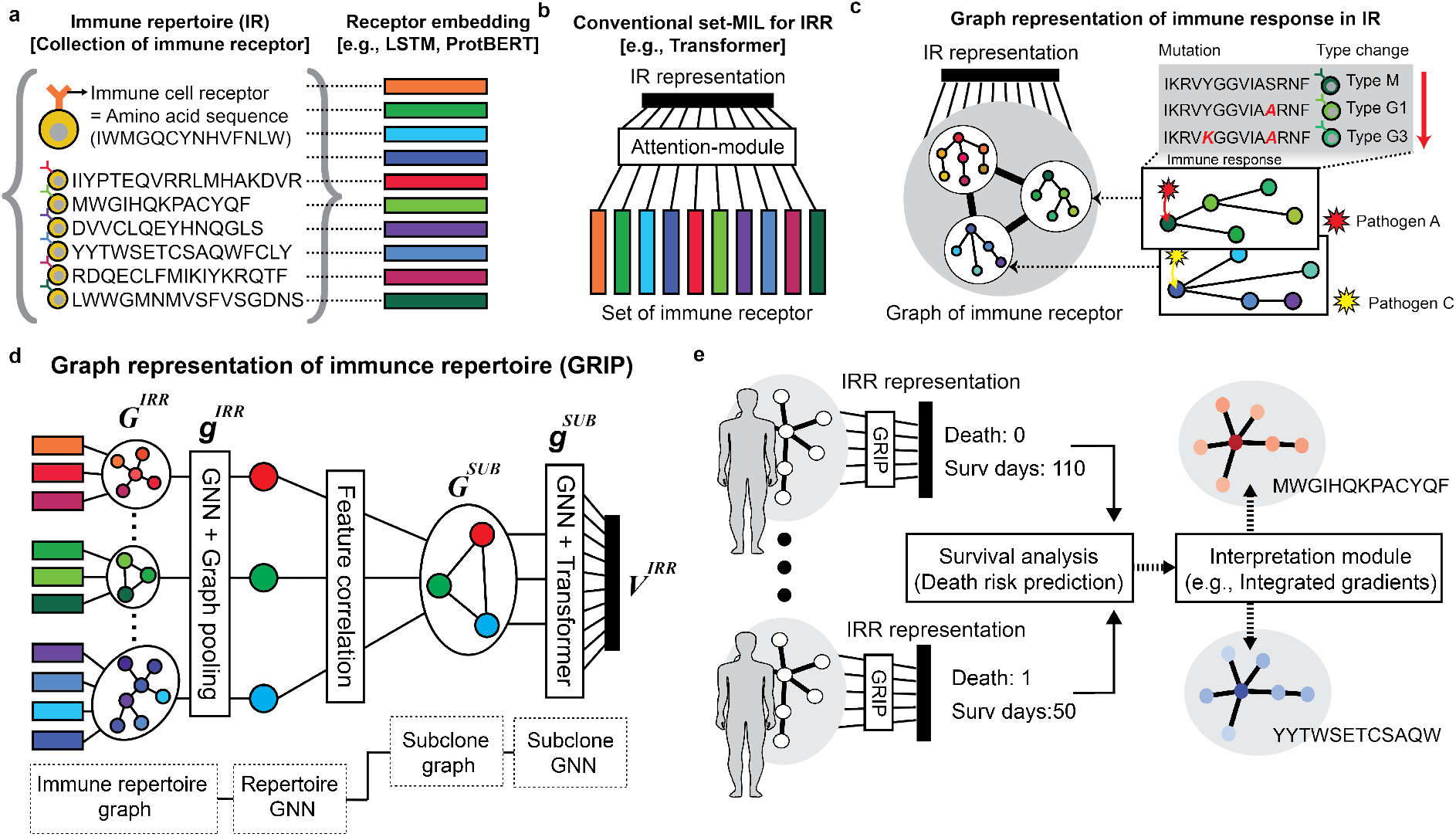
Overall schematic of a graph representation of immune receptor repertoire (GRIP). **a**, Immune repertoire is a collection of immune receptors, which is an amino acid sequence. We used the pre-trained protein language model to convert the immune receptor into an initial feature vector. **b**, Conventional set-level multiple instance learning (MIL) considers each immune receptor as an unrelated, unordered set. **c**, Immune cells introduce mutation and change isotype in the immune receptors when proliferating to defeat the pathogens as the immune response, and this is the lineage of the immune receptors. This lineage of an immune receptor is better represented as a graph. **d**, Structure of graph representation of immune repertoire (GRIP) that include cascaded graph neural network with graph pooling and transformer. **e**, Survival analysis and sequence-level interpretation of the cancer patients as an application of GRIP.

However, the immune repertoire is not a set-like data structure. Rather, inside the body, it forms a graph-like structure to defeat pathogens (Bashford-Rogers et al. 2013; Hozumi and Tonegawa 1976; Wu et al. 2015). Two types of immune receptors originate from T cells (T cell receptor, TCR) and B cells (B cell receptor, BCR). The BCR forms a more dynamic graph-like structure during the immune response. When B cells, at their pathogen-unexperienced state, encounter the pathogens that enter the body, they undergo proliferation and differentiation to optimize the BCR for the specific pathogens. The B cells introduce the mutations within the BCR (somatic hyper mutation, SHM) and change their isotype to efficiently defeat the pathogens (class switching recombination, CSR). The immune response sequentially and gradually optimizes the BCR. We can reconstruct the graph-like lineage of the BCR named subclone retrospectively by comparing the amino acid sequence of the BCR (Figure 1c). Thus, the immune repertoire is not a set-like structure but more similar to a graph-like structure with multiple lineages of BCR that react to pathogens in the body. Therefore, we need a framework that considers the graph property of immune repertoire data and analyzes the immune receptor sequence within them.

In this work, we introduced a framework that considers the graph-like properties of the immune response within the deep learning framework. We considered the immune repertoire data as a hierarchical graph structure and utilized a graph pooling strategy to efficiently analyze the immune receptor within the subclone. Specifically, we proposed a graph-based MIL model that maximizes the express power of a graph neural network (GNN) combined with a transformer model. The specific contributions of this study are as follows:

- We introduced a framework to represent the immune repertoire as a hierarchical graph structure, which reflects the subclone and repertoire-level immune response in the immune system.
- We proposed the graph representation learning of the immune repertoire (GRIP). GRIP is a cascaded graph neural network (GNN) that is combined graph pooling and transformer to learn the subclone-level and repertoire-level immune response within an immune repertoire.
- We demonstrated that GRIP predicts the survival probability of tumor patients using the immune repertoire of tumor tissue and yields interpretable results using the integrated gradients (IG) method. This proved that GRIP leverages the proposed graph structure of immune repertoires.

## Related Works

### MIL for immune repertoire analysis

Immune repertoire analysis is considered a MIL problem in several previous methods. DeepTCR introduced CNN and attention-module-based solution for TCR repertoire classification (Sidhom et al. 2021). DeepRC (Widrich et al. 2020) is a similar solution for immune repertoire classification. It uses LSTM for sequence embedding and an attention module for set-level MIL. DeepRC showed comparable results in classifying the normal and virus-infected immune repertoires. Additionally, a recent approach used the sequence embedding from a pre-trained antibody sequencing embedding model, AntiBERTa (Leem et al. 2022). They used the sequence features from AntiBERTa as the starting material. Subsequently, they conducted MIL using MLP, which revealed the proliferation characteristics of the immune repertoire through unsupervised clustering (Ruffolo, Gray, and Sulam 2021). Other approaches using k-mer motifs have also been introduced. However, these methods are limited in their ability to represent sequence features (Katayama and Kobayashi 2022). To the best of our knowledge, there are no other approaches that represent the immune repertoire as a graph and adapt a GNN for immune repertoire analysis.

### Graph-based analysis of the immune repertoire

The advantages of representing the immune repertoire as a graph have already been established in previous studies. Particularly, the graph characteristics of the immune repertoire help stratify the patients’ risk status against tumors. Previous researchers have used the statistical graph characteristics (e.g., Shannon’s entropy and evenness of graph) that demonstrated how strongly immune cells are differentiated, as well as the existence of certain major immune receptors (Yu et al. 2022). It is well known that the properties of the immune repertoire and prognosis of multiple tumor types, such as melanoma, bladder, kidney, and lung tumors, are related (Bolotin et al. 2017; Ferrall-Fairbanks et al. 2022; Harris et al. 2021; Isaeva et al. 2019; Petitprez et al. 2020; Zhan et al. 2022). In addition to its use for cancer prognosis, the graph characteristics of the immune repertoire help to classify the immune cell type and virus infection status (Miho et al. 2019; Pogorelyy et al. 2019). However, in all these studies, although the statistical graph characteristics were considered, the sequence-level features were not considered with them.

### Graph-based MIL

Although the graph-based MIL was not formally used term, a GNN naturally fell into the semi-supervised learning or graph-based MIL when we attempted to predict the graph-level features using the node and edge-level features of graph (Pal et al. 2022; Tu et al. 2019; Wu et al. 2022). The set-based MIL methods, such as DeepSet and AttentionMIL, are extensively studied in the computational pathology field where the giga-pixel whole slide image is used to predict the patient-level labels (Ilse, Tomczak, and Welling 2018; Li, Li, and Eliceiri 2021; Shao et al. 2021; Sharma et al. 2021; Yao et al. 2020; Zaheer et al. 2017). The recently introduced transformer is a powerful set-MIL model when it does not include the positional encoding, which is demonstrated in Set Transformer (Lee et al. 2019). The difference between set-based MIL and graph-based MIL is the inductive bias introduced to the instance that is used for message-passing during training. Therefore, if this inductive bias is helpful to the learning objective, graph-based MIL could be more powerful than set-based MIL. However, if it is not, the overall performance may be jeopardized. To compensate for this difficulty, we used a recently introduced GNN integrated with a transformer to maximize the express power of the GNN (Rámpašek et al. 2022; Wu et al. 2021).

## Methodology

### Problem formation

We regarded immune repertoire analysis as a graph-based MIL problem. An input immune repertoire graph G^IRR^ consists of N^IRR^ immune receptor *i*] ∈ *R*^d^ as a node. The number of immune receptors N^IRR^ can vary between immune repertoires. Let *A*^IRR^ ∈ *R*^N^^IRR^×^N^^IRR^ be an adjacency matrix describing the edge connection of the initial immune repertoire. The initial immune repertoire graph was pooled into the subclone graph G^SUB^. The subclone graph G^SUB^ is a bag of N^SUB^ subclone-level feature *s* ∈ *R^c^* as a node instance. Let *A*^SUB^ ∈ *R*^N^^SUB^ϗ^N^^SUB^ be an adjacency matrix describing the edge of the subclones. The subclone feature s is trained using the repertoire-level GNN g^IRR^ followed by mean-pooling based on the A^IRR^ that determines the N^SUB^. The immune repertoire representation vector *V*^IRR^ ∈ *R*^N^ is obtained using the subclone-level GNN g^SUB^ followed by a subclone-level transformer. Overall, we attempted to solve the graph-based MIL problem by learning a mapping g^IRR^:G^IRR^→G^SUB^,g^SUB^:G^SUB^→V^IRR^ where V^IRR^ is used to predict the graph-level label Y.

### Overview

We represented the immune repertoire as two different levels of the graph, which are the repertoire-level graph and the subclone-level graph. We proposed GRIP, a cascaded GNN combined with graph pooling and a transformer model to efficiently represent the immune response within the immune repertoire (Figure 1d). We attempted to predict the survival probability of cancer patients using the GRIP (Figure 1e).

We added an interpretation module to analyze the systematic properties of the immune repertoire and the immune sequences that are important for patient prognosis. Using the interpretation results, we can suggest the immune receptor sequence as the target for the biomarker.

### Graph representation of the immune repertoire

#### Immune repertoire graph construction

We constructed a graph from an immune repertoire based on the differences in the amino acids of two immune receptors. We defined the distance between the immune receptors as the character-level difference between two immune receptors and normalized it based on each immune receptor’s length. An edge was created between the two receptors when the normalized distance is less than 0.5.

#### Immune receptor sequence and CSR embedding

We used the pre-trained protein-level BERT model (Prot-BERT) to extract the initial embedding feature from each immune receptor sequence and initialized the entire node feature using the last layer of the ProtBERT model (Elnaggar et al. 2021). We applied a cascaded 1D CNN, followed by two layers of bidirectional GRU. We embedded the isotype of BCR as the learnable feature. We also calculated the number of each BCR sequence in the entire repertoire (frequency).

Overall, we concatenated the sequence-level feature, isotype feature, and frequency feature as the initial node feature for the immune repertoire graph. In addition, we measured the isotype changing between the BCR and used is as the CSR property. Concatenating with the normalized amino acid difference feature and CSR feature, we used it as the edge feature.

#### Subclone embedding by graph pooling

The subsets of immune receptors have very similar sequences, and these are called subclones. Also, the CSR feature in the subclone is important to represent the immune response. Through a graph pooling strategy with the CSR feature as an edge feature, we embedded these subclone-level features efficiently. We treated the subclone as the graph with a normalized distance of less than 0.2 between them. We used the three-layer GNN (repertoire level GNN, g^IRR^) to embed the immune receptor and CSR features of G^IRR^ within the subclone and pooled the node inside each subclone into a single node that represented the subclone-level feature as graph G^SUB^. We used GINE as the backbone model for g^IRR^ (Hu et al. 2019).

#### Immune repertoire embedding

After the subclone-level embedding, we conducted the repertoire-level embedding using the graph G^SUB^, which consisted of subclone-level embedding nodes. To construct the graph from the subclone-level embedding, we measured the cosine correlation between each subclone using the sequence-level features embedded through ProtBERT. We mean-pooled the sequence-level feature from ProtBERT for each subclone. When the cosine similarity of two subclones was over 0.9, we created an adjacency matrix A^SUB^ by connecting the edge between the subclone-level nodes. We then conducted a three-layer GNN g^SUB^ to embed the repertoire features. We concatenated the features from each GNN layer and used them for the initial features of the transformers. We used a two-layer transformer without positional encoding. We used the additional 〈*CLS*〉 token that represents the entire repertoire as the result of the final transformer layer. A detailed description of the model structure is included in the appendix.

### Survival analysis

#### Cox regression integrated with GRIP

We used GRIP to predict the survival probability (risk) of patients with cancer. To efficiently predict the risk of each patient, we adapted the survival analysis method of the Cox regression algorithm into the GNN (Katzman et al. 2018). Using the last repertoire representation vector V^IRR^, we predicted the loglikelihood of the Cox regression loss as follows:

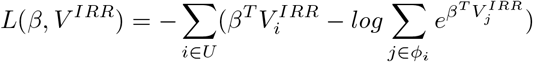

where, *β* ∈ *R* denotes the weights of the last MLP layer, U denotes the entire list of patients, and *φ* _i_ denotes the list of patients with shorter survival times than patient i. Moreover, censored patients are usually not included in cox regression loss calculation which makes it impossible to maximize the capability of entire datasets. We created an auxiliary loss that predicted the event status of the patient, which, in turn, used all the patients for training (Ren et al. 2019).

#### Interpretation

To derive prognostic features from the immune repertoire, it is important to interpret the results of GRIP. We adopted the IG method, which is known to be a reliable interpretation method for GNN (Sanchez-Lengeling et al. 2020; Sundararajan, Taly, and Yan 2017). In the survival analysis, the IG value of each node correlates with the risk of each node feature to the patient survival probability. Therefore, high and low IG values indicate high- and low-risk features, respectively.

## Experiments

### Experimental Setup

#### Dataset collection

We collected the bulk RNA-seq dataset of 14 different tumor types from the cancer genome atlas (TCGA) and extracted the immune receptor using TRUST, an algorithm for extracting immune receptors from bulk RNA-seq datasets (Song et al. 2021). Detailed dataset statistics are included in Table 1 and the appendix.

**Table 1:** Commands that must not be used

| Tumor type | HNSC | KIRP | LIHC | OV | PAAD | KIRC | COAD | LUSC | LUAD | BRCA | SKCM | BLCA | ESCA | READ | THCA | UCEC | STAD |
| --- | --- | --- | --- | --- | --- | --- | --- | --- | --- | --- | --- | --- | --- | --- | --- | --- | --- |
| # Patients | 358 | 130 | 165 | 286 | 127 | 351 | 326 | 372 | 409 | 806 | 364 | 249 | 147 | 124 | 258 | 203 | 295 |
| Avg #nodes | 395 | 227 | 90 | 466 | 435 | 207 | 254 | 926 | 804 | 312 | 334 | 222 | 802 | 263 | 535 | 35 | 1212 |
| Avg #edges | 7217 | 3993 | 399 | 6396 | 5067 | 1824 | 2074 | 16649 | 13464 | 3067 | 4087 | 1565 | 13191 | 3115 | 10698 | 53 | 24265 |
| Avg Degree | 29 | 35 | 8 | 27 | 23 | 17 | 16 | 35 | 33 | 19 | 24 | 14 | 32 | 23 | 39 | 2.98 | 40 |

#### Dataset preprocessing

We reconstructed the full-length BCR sequence using the aligned V and J gene information and the CDR3 sequence. We selected the immune receptor sequences with at least a length of five amino acids as the CDR3 sequence and immune repertoires that consisted of at least ten different immune receptors. To establish the link between the composition of immune cells within tumors, we used the inferred immune cell portion from previous studies as a global feature (Li et al. 2020). In addition, the prognostic power of immune receptors is strongly linked to DNA mutation status. Therefore, we collected the mutation status of 30 representative oncogenes as one-hot vectors from TCGA and embedded them in learnable global features (Gentles et al. 2015). A detailed description of dataset preprocessing is included in the appendix.

#### Baselines

We selected set-based methods (Attention-MIL, DeepSet, Set Transformer, and DeepRC) as the baseline for immune repertoire analysis. DeepSet is the permutation-invariant set-based deep learning method which is an adequate baseline for set-based MIL. Furthermore, although DeepRC uses the attention mechanism to solve the MIL problem, it used rather simplified attention mechanism. Therefore, we selected the Attention-MIL and Set Transformer models as an additional baseline for the advanced MIL model to solve the sophisticated immune repertoire datasets as the MIL problem. We used the same optimized hyperparameter for all methods.

#### Implementation details

We trained the GRIP using four RTX 3090. The learning rate was 0.001, with a weight decay and gradient clipping of 0.005 and 1.0, respectively. We used the polynomial decay learning rate scheduler with 20% of warmup steps for the entire training period. We conducted the hyperparameter screening and selected the hyperparameter that yielded the best results, as shown in Table 4.

**Table 2:** Commands that must not be used

| Tumor type | HNSC | KIRP | LIHC | OV | PAAD | KIRC | COAD | LUSC | LUAD | BRCA | SKCM | BLCA | ESCA | READ | THCA | UCEC | STAD | Total |
| --- | --- | --- | --- | --- | --- | --- | --- | --- | --- | --- | --- | --- | --- | --- | --- | --- | --- | --- |
| Att MIL | 0.6550 | 0.7651 | 0.5541 | 0.618 | 0.6689 | 0.6925 | 0.7212 | 0.5712 | 0.6284 | 0.6893 | 0.5685 | 0.6556 | 0.6250 | 0.7289 | 0.5927 | 0.7164 | 0.6143 | 0.6509 |
| DeepSet | 0.6527 | 0.6698 | 0.5135 | 0.5401 | 0.6591 | 0.6877 | 0.6731 | 0.5814 | 0.6609 | 0.726 | 0.6979 | 0.5519 | 0.6124 | 0.7721 | 0.6104 | 0.7313 | 0.6239 | 0.6450 |
| Set Trans | 0.6654 | 0.734 | 0.7083 | 0.6097 | 0.6578 | 0.7182 | **0.7404** | 0.6191 | 0.6513 | 0.7347 | 0.7128 | 0.6568 | 0.6125 | **0.7836** | 0.6223 | 0.806 | 0.6531 | 0.6874 |
| DeepRC | 0.6255 | 0.7617 | 0.6081 | 0.5353 | 0.6620 | 0.7156 | 0.6587 | 0.6084 | 0.6513 | 0.7574 | 0.7173 | 0.5477 | 0.6000 | 0.7500 | 0.5469 | 0.7313 | 0.6221 | 0.6529 |
| GRIP (ours) | **0.6822** | **0.7798** | **0.7235** | **0.6302** | **0.7072** | **0.7218** | 0.7260 | **0.6471** | **0.6667** | **0.7749** | **0.7619** | **0.6598** | **0.6750** | 0.7337 | **0.6698** | **0.8657** | **0.6871** | **0.7125** |

**Table 3:** Commands that must not be used

|  | HNSC | KIRP | LIHC | OV | PAAD | KIRC | COAD | LUAD | BRCA | SKCM | BLCA | ESCA | READ | THCA | UCEC | Total |
| --- | --- | --- | --- | --- | --- | --- | --- | --- | --- | --- | --- | --- | --- | --- | --- | --- |
| **GRIP (base)** | **0.6822** | **0.7798** | **0.7162** | **0.6302** | **0.7072** | 0.7218 | 0.726 | 0.6667 | 0.7749 | **0.7619** | **0.6598** | **0.675** | 0.7337 | 0.6698 | **0.8657** | 0.7181 |
| **w/o Graph pooling** | 0.6299 | 0.7312 | 0.7001 | 0.5877 | 0.6852 | 0.7213 | **0.7311** | **0.6771** | 0.7682 | 0.7412 | 0.6387 | 0.6611 | 0.7472 | **0.6714** | 0.8417 | 0.7022 |
| **w/o Repertoire GNN** | 0.6754 | 0.7429 | 0.7081 | 0.6112 | 0.6793 | **0.7355** | 0.7142 | 0.6384 | 0.7719 | 0.7429 | 0.6152 | 0.6529 | 0.7429 | 0.6223 | 0.8317 | 0.6990 |
| **w/o Subclone GNN** | 0.6678 | 0.7511 | 0.7155 | 0.6284 | 0.6826 | 0.7219 | 0.7075 | 0.6567 | **0.7811** | 0.7511 | 0.6459 | 0.6711 | 0.7511 | 0.6574 | 0.8565 | 0.7097 |
| **w/o Transformer** | 0.6331 | 0.7612 | 0.7135 | 0.6155 | 0.6787 | 0.7257 | 0.7198 | 0.6452 | 0.7532 | 0.7612 | 0.6523 | 0.6354 | **0.7612** | 0.6412 | 0.8444 | 0.7028 |

**Table 4:**
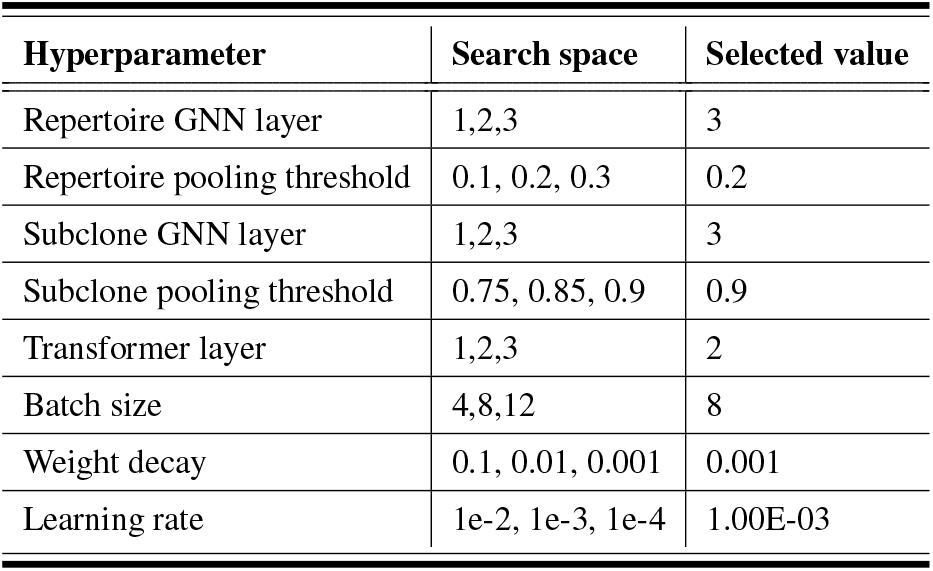
Commands that must not be used

## Results

### Performance Comparison

We compared the performance of GRIP with that of the baseline method for 17 different tumor types and 4,970 patients. The comparison results of the patient risk prediction are presented in Table 1. We measured the concordance index (C-index), which represents the accuracy of the survival analysis task. The C-index measures whether the predicted risks are ordered with respect to survival. For example, if one sample is predicted to have a higher risk than the other samples, the sample must have a shorter survival time. Thus, a 1.0 C-index indicated that all the predicted risk values were correctly ordered according to the survival time of each sample. GRIP outperformed the other set-based MIL methods for 15 out of 17 tumor types. Overall, GRIP outperformed the DeepRC and Set Transformer by 6% and 3%, respectively. In addition to the C-index, we also measured how well those methods stratify the at-risk patient group into high- or low-risk. For that, we divided the patients into high- and low-risk groups each patient group has a higher and lower predicted risk value than the median predicted risk value of entire patients. We plotted the results of the Kaplan-Meier analysis, which showed the patient survival rate over the entire timeline. We measured the area under the plot (AUP), which quantifies the difference between the high- and low-risk patient groups and the p-value using the paired ranked t-test. The higher AUP and lower p-value indicate the statistically well-stratified risk group. As shown in Figure. 2, GRIP stratified the patients into high- and low-risk groups more accurately than the Set Transformer in terms of both AUP and p-value. More results of patient stratification performance are included in the appendix.

**Figure 2:**
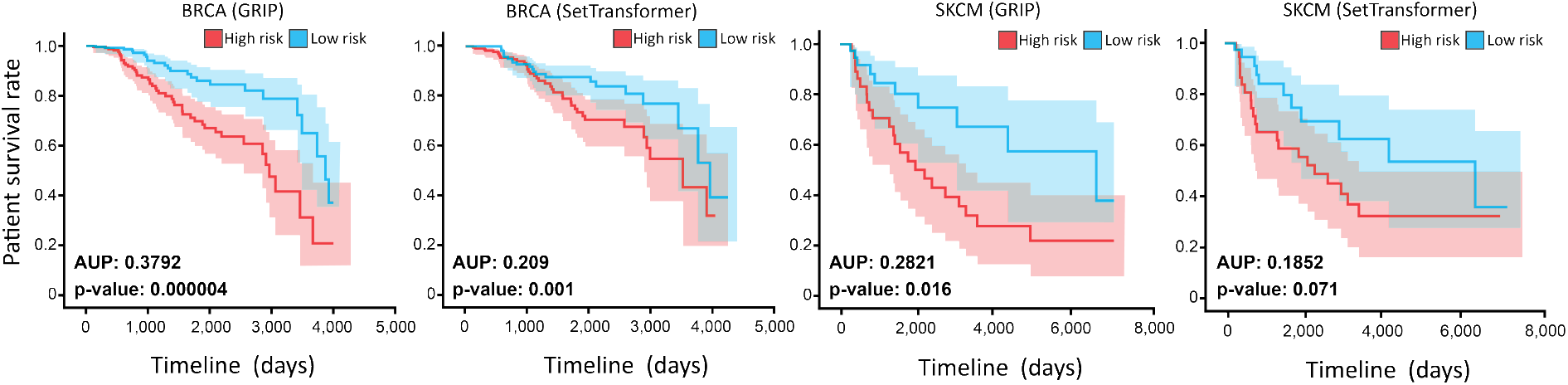
Kaplan-Meier analysis for patient stratification comparison of GRIP and Set Transformer

### Ablation Study

Through ablation studies, we evaluated the effectiveness of repertoire- and subclone-level GNN, graph-pooling, and transformer modules. The experimental results are listed in Table 3. The repertoire-level GNN is considered the graph of sequence in the subclone that shares most of the sequences but has minor differences in the mutated sequences. Furthermore, although they share similar sequences, they could have different isotypes indicating a status of the immune response. If we can pool the sequence consisting of subclones into a single subclone feature vector, we can efficiently process the following subclone-level learning. As expected, the repertoire GNN critically affected the performance and although graph pooling did not significantly affect the performance, we achieved comparable results with an 30% decrease in the number of nodes for subclone-level processing. Unlike the repertoire-level GNN, the sub-clone-level GNN considers the interaction between the subclone which is the overall immune status of an individual. We expected that the subclone-level GNN introduce the inductive bias that helps the model to better represent the immune repertoire than the set-based method that considers the whole pair of subclones independently. As expected, both the subclone GNN and transformer module affected the performance of GRIP, which implied that the subclone-level inductive bias we introduced through GNN provided helpful features to learn the immune response.

## Visualization and Interpretation

### Immune repertoire interpretation based on IG

Using the IG method, we measured the node-level IG score and used the mean of the entire node-level IG as the repertoire-level IG score. Also, we can obtain the attention value from the last layer of the transformer module. The IG value and attention value were used for the instance-level interpretation. We visualize the immune repertoire and colored the IG value as the node color. We defined the high/low IG subclone as a subgraph of nodes that shared a similar high- /low IG. We observed different amino properties between the high/low IG subclone (Figure 3a, appendix). Although GRIP and Set Transformer use the transformer module as the last layer, Set Transformer can’t consider the relationship between the immune receptor; class switching recombination (CSR). CSR is part of the immune response, which process of BCR optimization to better detect the pathogen that enters the body. To demonstrate the difference between the GRIP and Set Transformer, we first measured the attention value of each subclone and observed attention value of GRIP is sparser and more selective than Set Transformer (Figure 3b). Although further experiments are required, we speculate that GNN acts as the low pass filter prior to the transformer module in GRIP thus transformer more easily identifies the important features within the immune repertoire. Also, we measure the portion of CSR in the high/low IG group from GRIP and Set Transformer. In melanoma, G3 → G1 CSR is known to favorable prognosis marker for cancer patients (Petitprez et al. 2020). As expected, GRIP captures the appropriate CSR property that Set Transformer cannot consider. We found high IG group shows less G3 → G1 CSR than the low IG group in GRIP (Figure 3c). The G3 → G1 CSR is related to the mutation introduced in the BCR sequence during B cell proliferation and active communication of other immune cells within a tumor that helps to defeat the tumor cells (Hu et al. 2019). These were revealed through a statistical approach from previous studies. However, this is the first attempt to elucidate such results using deep learning. More results of high/low IG groups are included in the appendix.

**Figure 3:**
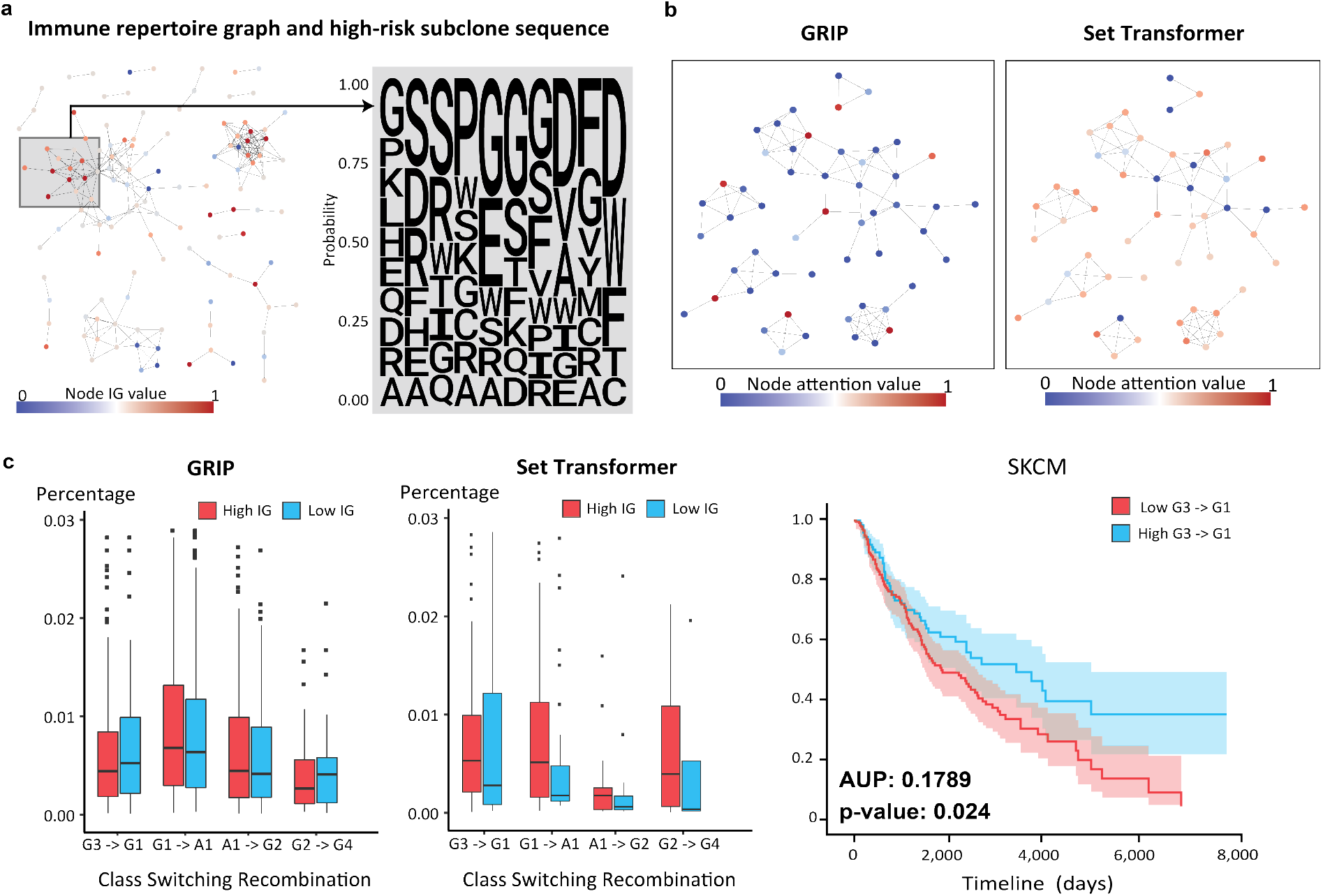
Visualization of immune repertoire and the comparison of GRIP and Set Transformer. **a**, Visualization of immune repertoire as a graph. Node color indicates the IG value. Red indicates a higher IG value which is more correlated with high risk. **b**, Comparison of attention value for each node between the GRIP and Set Transformer. **c**, Comparison of immune receptor properties between the high and low IG group of GRIP and Set Transformer.

## Conclusion

In this study, we developed GRIP, a novel method that represents the immune repertoire as a graph to better represent the immune response of the immune system. We introduced an immune repertoire graph construction method based on the sequence and feature similarities of each receptor and subclone. In addition, we introduced a cascaded GNN model structure followed by graph pooling and a transformer, which is an optimized model structure for analyzing the immune repertoire. We validated the performance of GRIP for the survival analysis of 5,000 number of cancer patients. It outperformed the set-based method, and we showed that GRIP considers the relationship between the immune cell receptor better than the set-based method using the interpretation module. Therefore, GRIP is the ideal method to consider both network structure and immune receptor sequence and to elucidate the connection between the network structure and immune receptor for immune response. In the future, we will further demonstrate the mechanism of GRIP that is different from the set-based method more thoroughly.

## Supporting information

Supplementary doc

## Acknowledgments

This work was supported by the Brain Korea 21 Plus Project in 2022, the Ministry of Science and ICT(MSIT) of the Republic of Korea and the National Research Foundation of Korea (NRF-2020R1A3B3079653).

## References

Bashford-Rogers, R. J. M.; Palser, A. L.; Huntly, B. J.; Rance, R.; Vassiliou, G. S.; Follows, G. A.; and Kellam, P. 2013. Network properties derived from deep sequencing of human B-cell receptor repertoires delineate B-cell populations. Genome Research, 23: 1874–1884.

Beshnova, D.; Ye, J.; Onabolu, O.; Moon, B.; Zheng, W.; Fu, Y.-X.; Brugarolas, J.; Lea, J.; and Li, B. 2020. De novo prediction of cancer-associated T cell receptors for noninvasive cancer detection. Science Translational Medicine, 12.

Bolotin, D. A.; Poslavsky, S.; Davydov, A. N.; Frenkel, F. E.; Fanchi, L.; Zolotareva, O. I.; Hemmers, S.; Putintseva, E. V.; Obraztsova, A. S.; Shugay, M.; et al. 2017. Antigen receptor repertoire profiling from RNA-seq data. Nature Biotechnology, 35: 908–911.

Davidsen, K.; Olson, B. J.; DeWitt, W. S.; Feng, J.; Harkins, E.; Bradley, P.; and Matsen, F. A. 2019. Deep generative models for T cell receptor protein sequences. eLife, 8: e46935.

Elnaggar, A.; Heinzinger, M.; Dallago, C.; Rehawi, G.; Wang, Y.; Jones, L.; Gibbs, T.; Feher, T.; Angerer, C.; Steinegger, M.; et al. 2021. ProtTrans: Towards Cracking the Language of Lifes Code Through Self-Supervised Deep Learning and High Performance Computing. IEEE Transactions on Pattern Analysis and Machine Intelligence, PP: 1.

Emerson, R. O.; DeWitt, W. S.; Vignali, M.; Gravley, J.; Hu, J. K.; Osborne, E. J.; Desmarais, C.; Klinger, M.; Carlson, C. S.; Hansen, J. A.; et al. 2017. Immunosequencing identifies signatures of cytomegalovirus exposure history and HLA-mediated effects on the T cell repertoire. Nature Genetics, 49: 659–665.

Ferrall-Fairbanks, M. C.; Chakiryan, N.; Chobrutskiy, B. I.; Kim, Y.; Teer, J. K.; Berglund, A.; Mulé, J. J.; Fournier, M.; Siegel, E. M.; Dhillon, J.; et al. 2022. Quantification of T-and B-cell immune receptor distribution diversity characterizes immune cell infiltration and lymphocyte heterogeneity in clear cell renal cell carcinomaClinical relevance of adaptive immune cell heterogeneity. Cancer research.

Gentles, A. J.; Newman, A. M.; Liu, C. L.; Bratman, S. V.; Feng, W.; Kim, D.; Nair, V. S.; Xu, Y.; Khuong, A.; Hoang, C. D.; et al. 2015. The prognostic landscape of genes and infiltrating immune cells across human cancers. Nature Medicine, 21: 938–945.

Greiff, V.; Weber, C. R.; Palme, J.; Bodenhofer, U.; Miho, E.; Menzel, U.; and Reddy, S. T. 2017. Learning the High Dimensional Immunogenomic Features That Predict Public and Private Antibody Repertoires. The Journal of Immunology, 199: 2985–2997.

Harris, R. J.; Cheung, A.; Ng, J. C. F.; Laddach, R.; Chenoweth, A. M.; Crescioli, S.; Fittall, M.; Dominguez-Rodriguez, D.; Roberts, J.; Levi, D.; et al. 2021. Tumor Infiltrating B Lymphocyte Profiling Identifies IgG-Biased, Clonally Expanded Prognostic Phenotypes in Triple Negative Breast Cancer. Cancer Research, 81: 4290–4304.

Hozumi, N.; and Tonegawa, S. 1976. Evidence for somatic rearrangement of immunoglobulin genes coding for variable and constant regions. Proceedings of the National Academy of Sciences, 73: 3628–3632.

Hu, X.; Zhang, J.; Wang, J.; Fu, J.; Li, T.; Zheng, X.; Wang, B.; Gu, S.; Jiang, P.; Fan, J.; et al. 2019. Landscape of B cell immunity and related immune evasion in human cancers. Nature Genetics, 51: 560–567.

Ilse, M.; Tomczak, J.; and Welling, M. 2018. Attention-based deep multiple instance learning. In International conference on machine learning, 2127–2136. PMLR.

Isaeva, O. I.; Sharonov, G. V.; Serebrovskaya, E. O.; Turchaninova, M. A.; Zaretsky, A. R.; Shugay, M.; and Chudakov, D. M. 2019. Intratumoral immunoglobulin isotypes predict survival in lung adenocarcinoma subtypes. Journal for ImmunoTherapy of Cancer, 7: 279.

Jin, W.; Wohlwend, J.; Barzilay, R.; and Jaakkola, T. S. 2021. Iterative Refinement Graph Neural Network for Antibody Sequence-Structure Co-design. In International Conference on Learning Representations.

Katayama, Y.; and Kobayashi, T. J. 2022. Comparative Study of Repertoire Classification Methods Reveals Data Efficiency of k -mer Feature Extraction. Frontiers in Immunology, 13: 797640.

Katzman, J. L.; Shaham, U.; Cloninger, A.; Bates, J.; Jiang, T.; and Kluger, Y. 2018. DeepSurv: personalized treatment recommender system using a Cox proportional hazards deep neural network. BMC Medical Research Methodology, 18: 24.

Lee, J.; Lee, Y.; Kim, J.; Kosiorek, A.; Choi, S.; and Teh, Y. W. 2019. Set transformer: A framework for attention-based permutation-invariant neural networks. In International conference on machine learning, 3744–3753. PMLR

Leem, J.; Mitchell, L. S.; Farmery, J. H. R.; Barton, J.; and Galson, J. D. 2022. Deciphering the language of antibodies using self-supervised learning. Patterns, 3: 100513.

Li, B.; Li, Y.; and Eliceiri, K. W. 2021. Dual-stream multiple instance learning network for whole slide image classification with self-supervised contrastive learning. In Proceedings of the IEEE/CVF conference on computer vision and pattern recognition, 14318–14328.

Li, T.; Fu, J.; Zeng, Z.; Cohen, D.; Li, J.; Chen, Q.; Li, B.; and Liu, X. S. 2020. TIMER2. 0 for analysis of tumorinfiltrating immune cells. Nucleic acids research, 48(W1): W509–W514.

Mason, D. M.; Friedensohn, S.; Weber, C. R.; Jordi, C.; Wagner, B.; Meng, S. M.; Ehling, R. A.; Bonati, L.; Dahinden, J.; Gainza, P.; et al. 2021. Optimization of therapeutic antibodies by predicting antigen specificity from antibody sequence via deep learning. Nature Biomedical Engineering, 5: 600–612.

Miho, E.; Roškar, R.; Greiff, V.; and Reddy, S. T. 2019. Large-scale network analysis reveals the sequence space architecture of antibody repertoires. Nature Communications, 10: 1321.

Mora, T.; and Walczak, A. M. 2019. How many different clonotypes do immune repertoires contain? Current Opinion in Systems Biology, 18: 104–110.

Pal, S.; Valkanas, A.; Regol, F.; and Coates, M. 2022. Bag Graph: Multiple Instance Learning using Bayesian Graph Neural Networks. In Proc. AAAI Conf. on Artificial Intelligence.

Petitprez, F.; de Reyniès, A.; Keung, E. Z.; Chen, T. W.-W.; Sun, C.-M.; Calderaro, J.; Jeng, Y.-M.; Hsiao, L.-P.; Lacroix, L.; Bougoüin, A.; et al. 2020. B cells are associated with survival and immunotherapy response in sarcoma. Nature, 577: 1–5.

Pogorelyy, M. V.; Minervina, A. A.; Shugay, M.; Chudakov, D. M.; Lebedev, Y. B.; Mora, T.; and Walczak, A. M. 2019. Detecting T cell receptors involved in immune responses from single repertoire snapshots. PLOS Biology, 17: e3000314.

Rampášek, L.; Galkin, M.; Dwivedi, V. P.; Luu, A. T.; Wolf, G.; and Beaini, D. 2022. Recipe for a General, Powerful, Scalable Graph Transformer. arXiv preprint arXiv:2205.12454.

Ren, K.; Qin, J.; Zheng, L.; Yang, Z.; Zhang, W.; Qiu, L.; and Yu, Y. 2019. Deep recurrent survival analysis. In Proceedings of the AAAI Conference on Artificial Intelligence, volume 33, 4798–4805.

Ruffolo, J. A.; Gray, J. J.; and Sulam, J. 2021. Deciphering antibody affinity maturation with language models and weakly supervised learning. arXiv preprint arXiv:2112.07782.

Sanchez-Lengeling, B.; Wei, J.; Lee, B.; Reif, E.; Wang, P.; Qian, W.; McCloskey, K.; Colwell, L.; and Wiltschko, A. 2020. Evaluating attribution for graph neural networks. Advances in neural information processing systems, 33: 5898–5910.

Shao, Z.; Bian, H.; Chen, Y.; Wang, Y.; Zhang, J.; Ji, X.; et al. 2021. Transmil: Transformer based correlated multiple instance learning for whole slide image classification. Advances in Neural Information Processing Systems, 34: 2136–2147.

Sharma, Y.; Shrivastava, A.; Ehsan, L.; Moskaluk, C. A.; Syed, S.; and Brown, D. 2021. Cluster-to-conquer: A framework for end-to-end multi-instance learning for whole slide image classification. In Medical Imaging with Deep Learning, 682–698. PMLR.

Sidhom, J.-W.; Larman, H. B.; Pardoll, D. M.; and Baras, A. S. 2021. DeepTCR is a deep learning framework for revealing sequence concepts within T-cell repertoires. Nature Communications, 12: 1605.

Slamon, D.; Eiermann, W.; Robert, N.; Pienkowski, T.; Martin, M.; Press, M.; Mackey, J.; Glaspy, J.; Chan, A.; Pawlicki, M.; et al. 2011. Adjuvant Trastuzumab in HER2-Positive Breast Cancer. The New England Journal of Medicine, 365: 1273–1283.

Song, L.; Cohen, D.; Ouyang, Z.; Cao, Y.; Hu, X.; and Liu, X. S. 2021. TRUST4: immune repertoire reconstruction from bulk and single-cell RNA-seq data. Nature Methods, 18: 627–630.

Sundararajan, M.; Taly, A.; and Yan, Q. 2017. Axiomatic attribution for deep networks. In International conference on machine learning, 3319–3328. PMLR.

Thorsson, V.; Gibbs, D. L.; Brown, S. D.; Wolf, D.; Bortone, D. S.; Yang, T.-H. O.; Porta-Pardo, E.; Gao, G. F.; Plaisier, C. L.; Eddy, J. A.; et al. 2018. The Immune Landscape of Cancer. Immunity, 48: 812–830.e14.

Tu, M.; Huang, J.; He, X.; and Zhou, B. 2019. Multiple instance learning with graph neural networks. arXiv preprint arXiv:1906.04881.

Widrich, M.; Schäfl, B.; Pavlović, M.; Ramsauer, H.; Gruber, L.; Holzleitner, M.; Brandstetter, J.; Sandve, G. K.; Greiff, V.; Hochreiter, S.; et al. 2020. Modern Hopfield Networks and Attention for Immune Repertoire Classification. bioRxiv, 2020.04.12.038158.

Wu, F.; Liu, P.; Fu, B.; and Ye, F. 2022. DeepGCNMIL: Multi-head Attention Guided Multi-Instance Learning Approach for Whole-Slide Images Survival Analysis Using Graph Convolutional Networks. In 2022 14th International Conference on Machine Learning and Computing (ICMLC), 67–73.

Wu, X.; Zhang, Z.; Schramm, C. A.; Joyce, M. G.; Kwon, Y. D.; Zhou, T.; Sheng, Z.; Zhang, B.; O’Dell, S.; McKee, K.; et al. 2015. Maturation and Diversity of the VRC01-Antibody Lineage over 15 Years of Chronic HIV-1 Infection. Cell, 161: 470–485.

Wu, Z.; Jain, P.; Wright, M.; Mirhoseini, A.; Gonzalez, J. E.; and Stoica, I. 2021. Representing long-range context for graph neural networks with global attention. Advances in Neural Information Processing Systems, 34: 13266–13279.

Yao, J.; Zhu, X.; Jonnagaddala, J.; Hawkins, N.; and Huang, J. 2020. Whole slide images based cancer survival prediction using attention guided deep multiple instance learning networks. Medical Image Analysis, 65: 101789.

Yu, K.; Ravoor, A.; Malats, N.; Pineda, S.; and Sirota, M. 2022. A Pan-Cancer Analysis of Tumor-Infiltrating B Cell Repertoires. Frontiers in Immunology, 12: 790119.

Zaheer, M.; Kottur, S.; Ravanbakhsh, S.; Poczos, B.; Salakhutdinov, R. R.; and Smola, A. J. 2017. Deep sets. Advances in neural information processing systems, 30.

Zhan, Y.; Guan, X.; Zhang, Y.; Zhu, Z.; Shi, A.; and Fan, Z. 2022. Identification of an immune-related gene pair signature in breast cancer. Translational Cancer Research, 0: 0.

