## Supplementary doc for "GRIP: Graph Representation of Immune Repertoire Using Graph Neural Network and Transformer"

**Anonymous Submission**

### Dataset preprocessing

**Immune receptor sequence extraction and preprocessing with ProtBERT feature vector**

We download the RNA-seq BAM file dataset for the entire 14 different tumor types in the cancer genome atlas (TCGA) through gdc portal (https://portal.gdc.cancer.gov/). We rule out the normal sample included in each dataset. And then, we extracted the immune receptor from the bulk RNA-seq BAM file. Recently, several methods can be used for immune receptor extraction, such as MiXCR, TRUST, and VDJer. We want to create a meaningful immune repertoire graph from the extracted sequence, so we want to gather as many immune receptors as possible but avoid false positive ones from TCGA dataset. So, we choose the TRUST method that is already used for TCGA dataset analysis and capable of extracting as many immune receptors as possible in a highly sensitive manner.

Owing to the limitation of sensitivity in bulk RNA sequencing, we rarely recover the full portion of the B cell receptor (BCR) sequence and T cell receptor (TCR) sequence from TCGA dataset. However, we can reconstruct the partially obtained BCR using the special characteristics of BCR. The BCR has a high mutation region called the complementarity-determining region (CDR) and a relatively conserved germline region (V, J gene). Because the germline region is conserved within BCR, we can easily identify the V and J gene types from the few sequences of recovered BCR sequences. Also, there are three CDR portions in BCR such as CDR1, CDR2, and CDR3. Among them, CDR3 is known to be the most important region that changed the binding property of BCR. Therefore, we only selected the BCR where both V and J gene types are identified, and the full CDR3 sequence is recovered. We reconstruct the full BCR sequence using the archived V, J gene sequence matched with identified V, J gene types.

The average length of the reconstructed BCR sequence is ~150 amino acid sequence, and immune repertoire sequence data usually consist of 1000 ~ 10000 number of BCR, which is not easy to be learned through the GPU. Therefore, we compress sequence features by mean-pooling the ProtBERT features from the V gene and J gene, which is a highly conserved region. We mean-pooled every five amino acid sequence into one sequence feature chunk, thus making ~105 amino acid V gene into ~21 V gene feature chunk and ~15 amino acid J gene into ~3 J gene feature chunk. As a result, we concatenate the V gene feature chunk, naïve CDR3 sequence feature extracted using ProtBERT, and J gene feature chunk as the feature vector for each immune receptor. Overall, we prepared the immune repertoire data for each individual cancer patient that consist of a compressed BCR sequence feature vector.

**Summarized immune repertoire and subclone graph property**

Each tumor type has a different number of BCR and different sequence-level similarities to each other. We summarized immune repertoire and subclone graph property that possibly affect the graph-pooling and performance of GNN in Table 1.

### Model structure

**Initial node feature embedding**

We applied the two layers of 1D CNN (1024 x 128, 128 x 128) followed by two layers of weight normalized bidirectional GRU (128 x 128) for BCR sequence embedding. We used the GELU for the activation function of 1D CNN (stride: 1, kernel size: 3) with Max-pooling (kernel size: 3). We embedded ten different isotypes into learnable features (1 x 20). We calculated frequency as the ratio of specific BCR among the entire immune repertoire and quantized the frequency feature into 0 – 10 numbers. And then, we created lookup tables for frequency features as learnable feature vectors. We concatenated the flattened sequence features from GRU, isotype, and frequency features as a single vector. We applied the two-layer MLP and used the features from the last layer of MLP as the node feature.

| | | HNSC | KIRP | LIHC | OV | PAAD | KIRC | COAD | LUSC | LUAD | CESC | BRCA | SKCM | BLCA | ESCA | READ | THCA | UCEC | STAD | Total |
|---|---|---|---|---|---|---|---|---|---|---|---|---|---|---|---|---|---|---|---|---|
| # Patients | | 358 | 130 | 165 | 286 | 127 | 351 | 326 | 372 | 409 | 231 | 806 | 364 | 249 | 147 | 124 | 258 | 203 | 295 | 5201 |
| Repertoire | Avg #nodes | 2.67 | 2.50 | 3.09 | 3.48 | 2.67 | 2.67 | 2.60 | 2.63 | 2.77 | 2.61 | 2.78 | 3.00 | 2.62 | 2.97 | 2.59 | 2.93 | 2.36 | 3.00 | 2.77 |
| | Avg #edges | 1.40 | 1.26 | 1.57 | 2.31 | 1.35 | 1.28 | 1.30 | 1.34 | 1.48 | 1.32 | 1.34 | 1.67 | 1.29 | 1.92 | 1.33 | 1.47 | 1.07 | 1.87 | 1.48 |
| Subclone | Avg #nodes | 395 | 227 | 90 | 466 | 435 | 207 | 254 | 926 | 804 | 306 | 312 | 334 | 222 | 802 | 263 | 535 | 35 | 1212 | 434.72 |
| | Avg #edges | 7217 | 3993 | 399 | 6396 | 5067 | 1824 | 2074 | 16649 | 13464 | 3151 | 3067 | 4087 | 1565 | 13191 | 3115 | 10698 | 53 | 24265 | 6681.94 |
| | Avg Degree | 29 | 35 | 8 | 27 | 23 | 17 | 16 | 35 | 33 | 20 | 19 | 24 | 14 | 32 | 23 | 39 | 2.98 | 40 | 24.28 |

Table 1: Graph characteristics of repertoire-level and subclone-level graph

**Initial edge feature embedding**

We embedded the class switching recombination (CSR) as learnable features and quantized the normalized sequence difference into 0 – 10 numbers. Like the frequency feature, we created lookup tables for sequence difference as a learnable feature vector. We concatenated the CSR and distance feature into one single vector and applied the two-layer MLP.

**Initial global feature embedding**

We select 29 oncogenes that are well known for their effect on cancer prognosis and relation to the tumor microenvironment. The list of oncogenes as follows (ATM, POLE, TGFB2, 16p, BRAF, MLH1, CTCF, PIK3R1, ARID1A, PTEN, PIK3CA, 20q, APC, TP53, NRAS, KRAS, MGMT, CDKN2A, EGFR, PDGFRA, IDH1, DNMT3A, NPM1, FLT3, BAP1, SETD2, GSTP1, PBRM1, VHL). We created the one-hot encoding table using the mutation information from TCGA dataset and embedded the mutation information using the lookup table as the learnable feature. Also, we include a feature for the cell type portion and activation status within tumor mass for each patient. The included immune cell features are as follows (leukocyte, stromal, infiltrating lymphocyte fraction, Th1, Th2, Th17, Memory B cell, Naïve B cell, Dendritic cell, Eosinophils, Macrophage, Mast cell, Monocyte, Neutrophil, NK cell, Plasma cell, T cell CD4, T cell CD8 ). We used the raw float number as the input feature. Overall, we concatenate the oncogene mutation feature and cell type portion feature as a single feature vector.

**Repertoire-level GNN followed by graph pooling**

With the initialized node, edge feature, and repertoire-level adjacency matrix calculated based on the amino acid difference between two immune receptors, we conducted three-layer of GNN (128 x 128). We used the GINE as the backbone model to use the node and edge features simultaneously. We add the residual connection between each GNN layer and concatenate the feature that comes from each layer of GNN. We mean-pool the results of repertoire-level GNN that included in the same subclone (< 0.2 amino acid difference).

**Subclone-level GNN followed by transformer**

With the mean-pooled node feature, global feature, and subclone-level adjacency matrix calculated using the sequence-level ProtBERT feature, we conducted three-layer of GNN (128 x 128). We used the GIN as the backbone model and added the residual connection between each GNN layer. We concatenate the input node feature and feature that come from each layer of GNN. We used the embedded features from subclone-level GNN as the input feature for three-layer of a transformer (512 x 512). We used the transformer that has four, two, and one attention head for each transformer layer. Also, we include the <CLS> token for the transformer as the learnable feature that represents the entire subclone-graph as the single embedding feature.

**Survival probability prediction**

We concatenate the <CLS> token feature from the last layer of a transformer, mean-pooled features from the last layer of a transformer, and the output of the subclone-level GNN feature as the single vector to predict the survival risk of patients. We applied two-layer MLP for the flattened feature and used the results as the input to calculate the cox regression loss.

**Auxiliary loss for survival loss and repertoire-level GNN**

We introduce the auxiliary loss to maximize the efficiency of survival analysis and prevent the overfitting issue in GNN. We applied two-layer MLP for the flattened feature, which is the same feature used for cox regression loss, and used the results to predict the survival status of each patient. Also, we replace the 0.5% input node feature with random noise and predict the replaced node feature with the aggregated neighborhood features from repertoire-level GNN.

### Patient stratification results

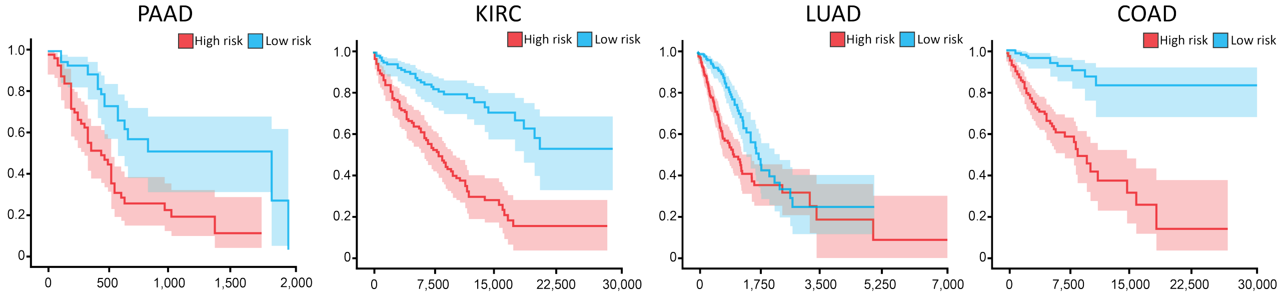


Figure 1: Patient stratification results using the predicted risk from GRIP

### Visualization and interpretation

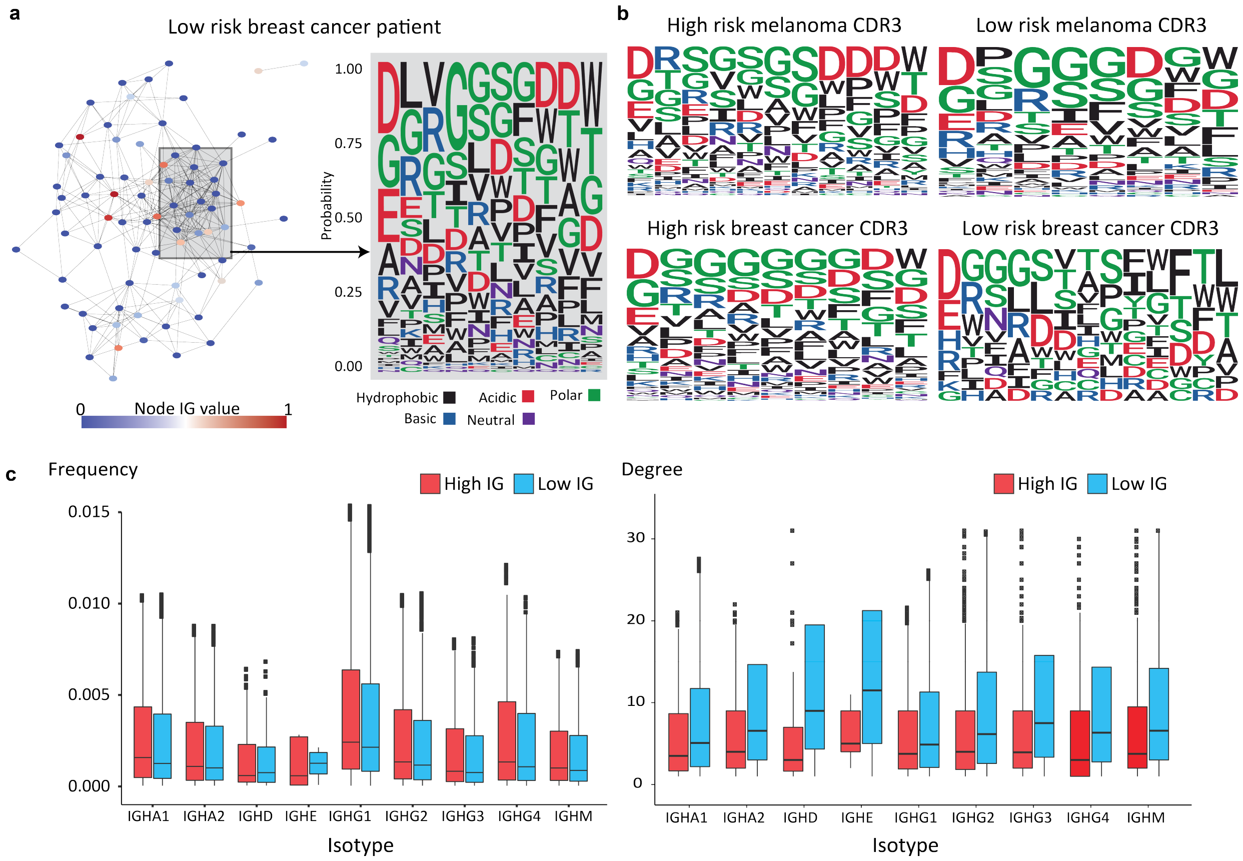


Figure 2: Immune repertoire visualization and difference between the high and low IG group

We define the subclone as the connected subgraph that has a similar IG value and obtain the sequence-level property from the low IG and high IG patient groups (Figure 2a). Although further both data and bioassay-based experiments are required, as shown in Figure 2b, high risk (high IG) and low risk (low IG) CDR3 sequences in each tumor type have different electro-physiological properties. We can measure the different frequency characteristics of which IGHD isotype more frequently exist in the low IG group than in the high IG group. In addition, the low IG group has a higher degree than the high IG group, which shows that the low IG group has a more active immune cell proliferation status.
